## Supplementary figures and images for "Mapping responses to focal injections of bicuculline in the lateral parafacial region identifies core regions for maximal generation of active expiration"

### supplemental 1 gif

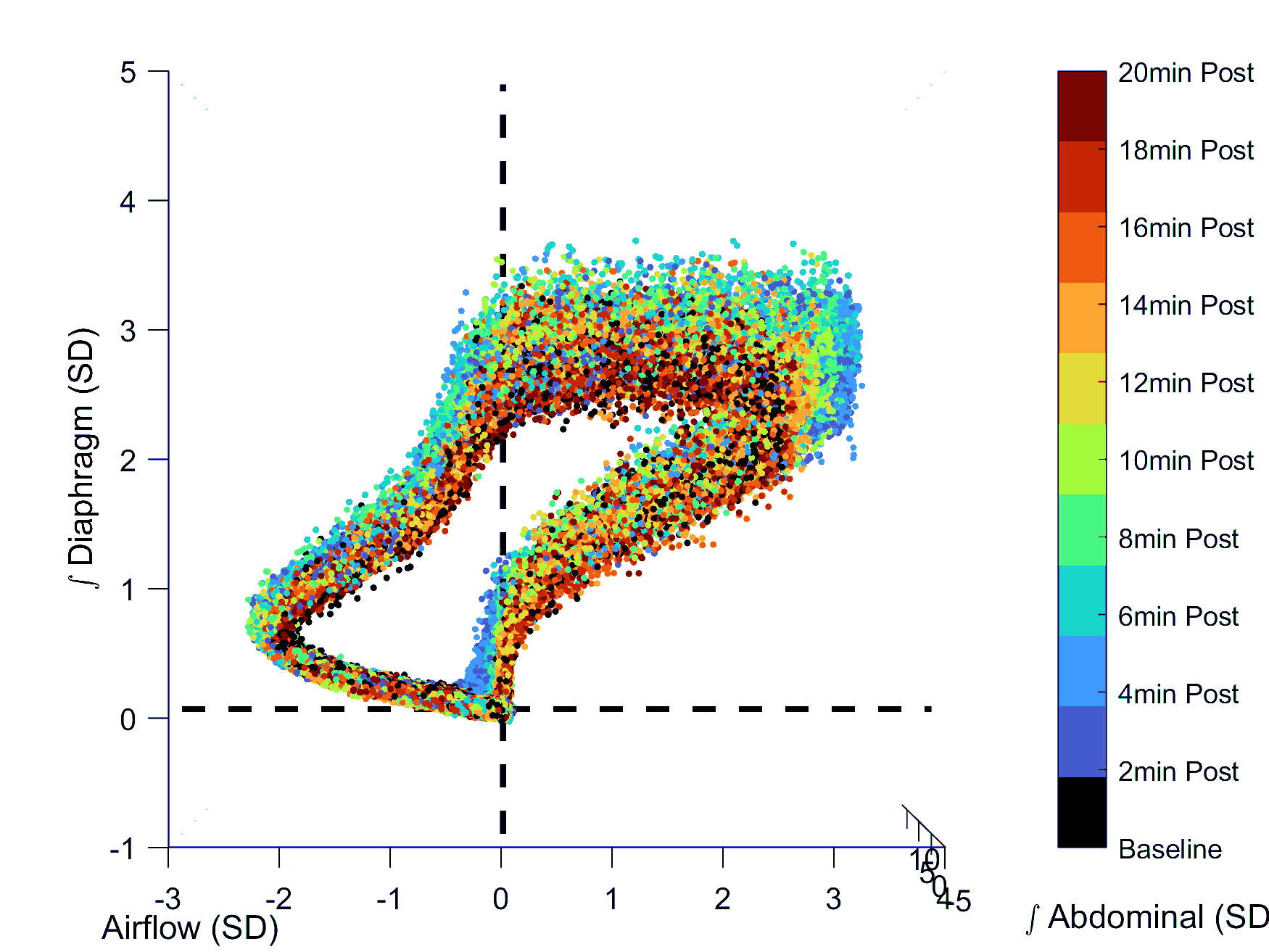

### supplemental 2 gif

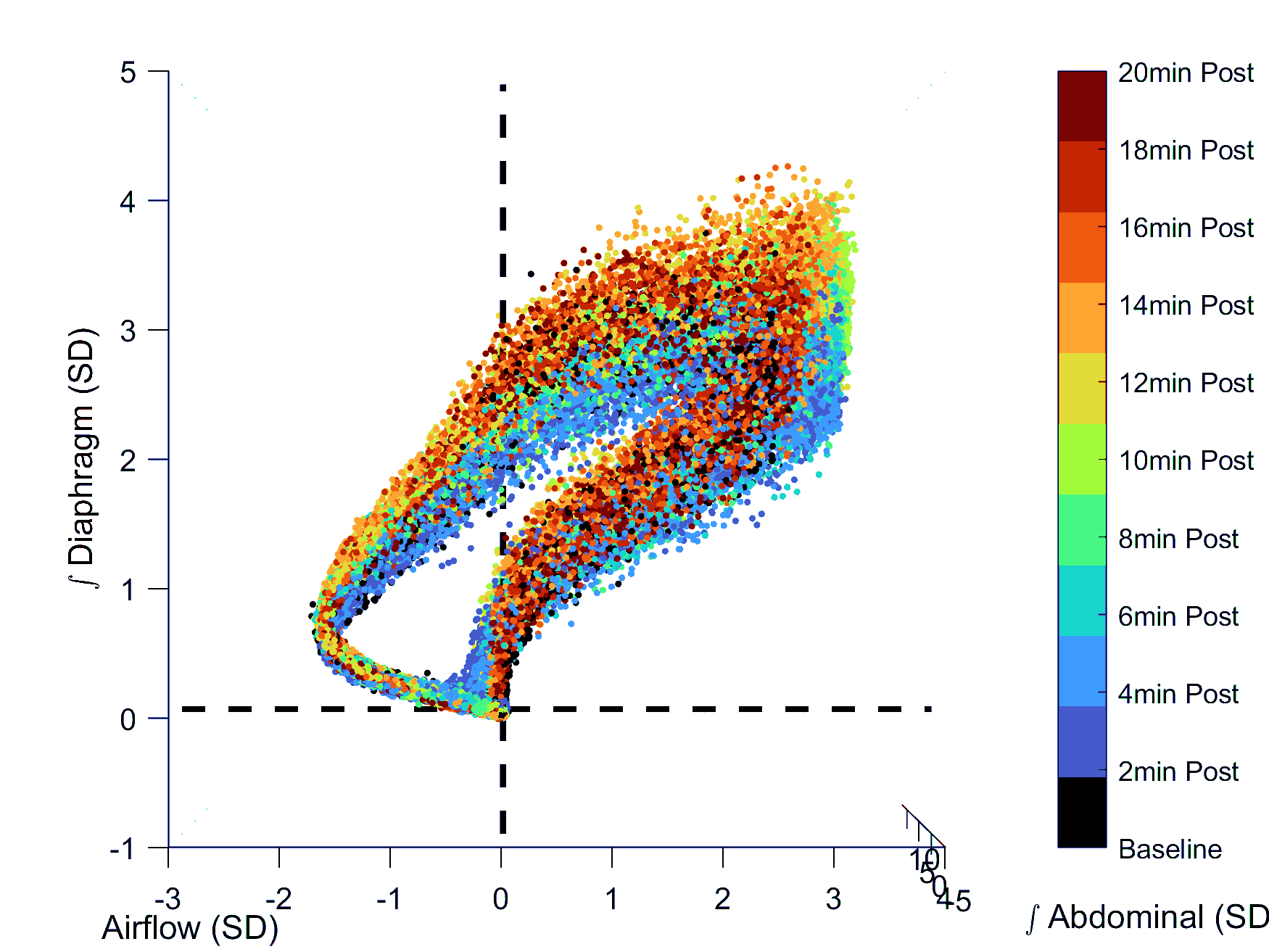

### supplemental 3 gif

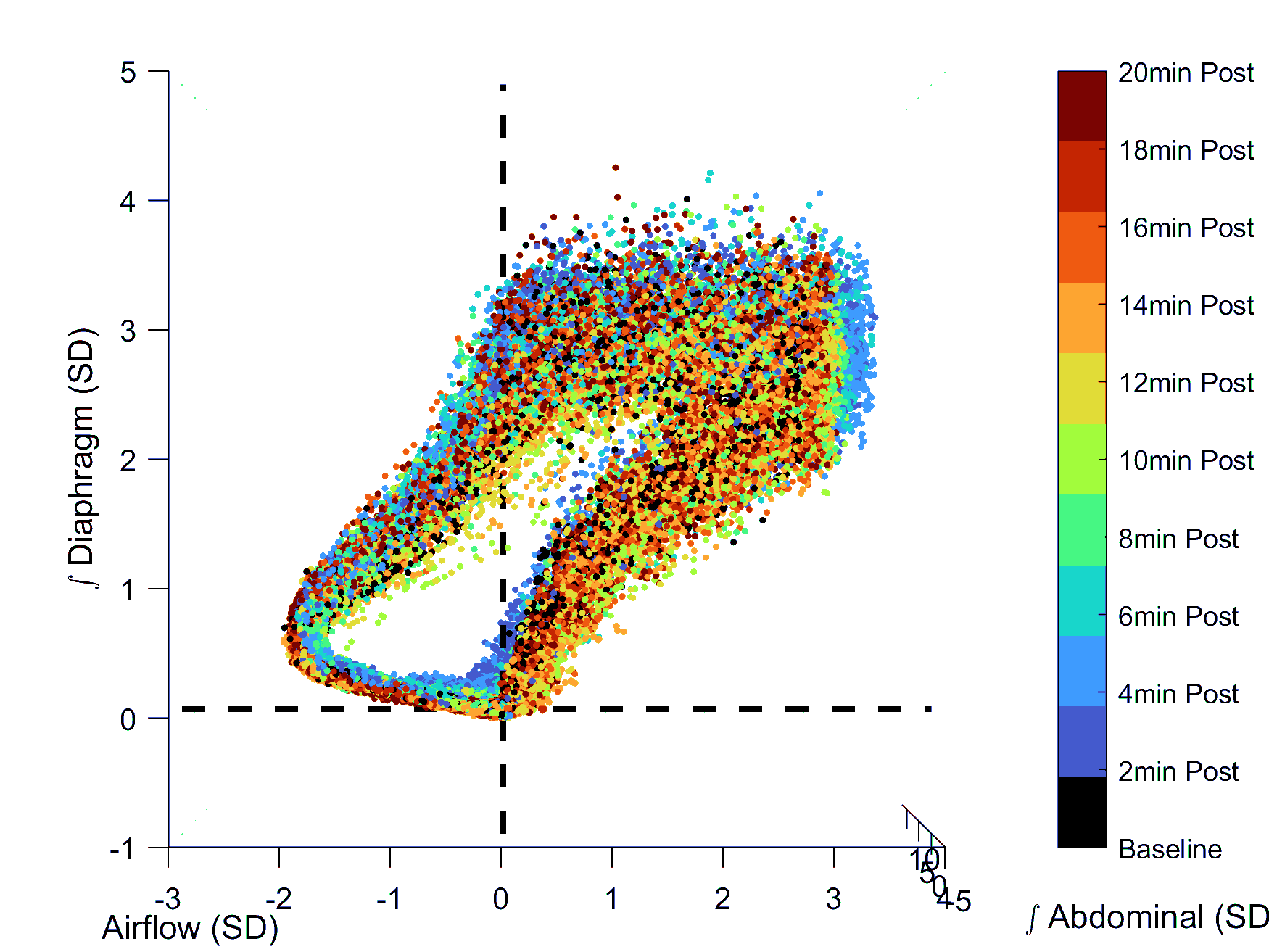

### supplemental 4 gif

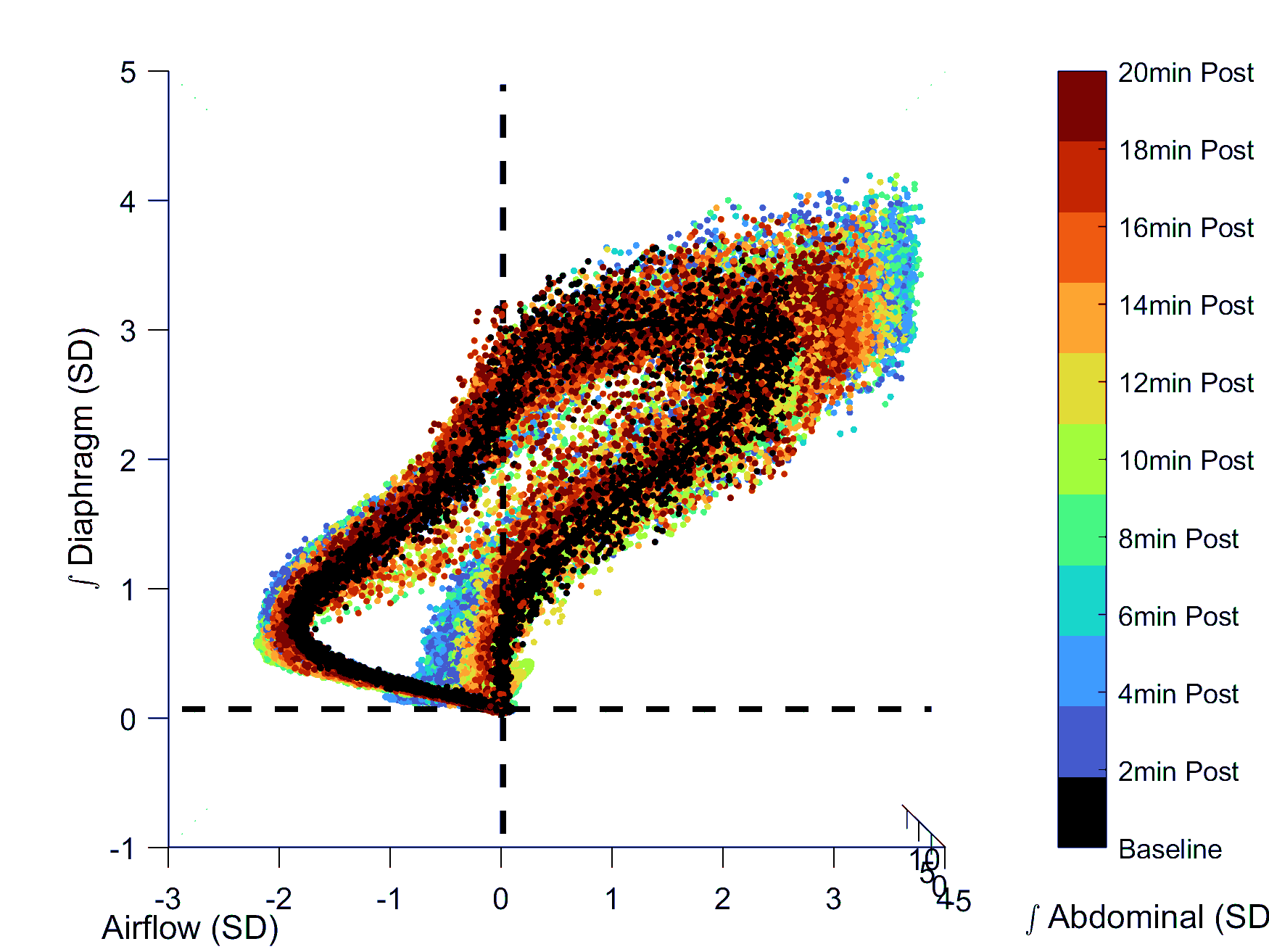

### supplemental 5 gif

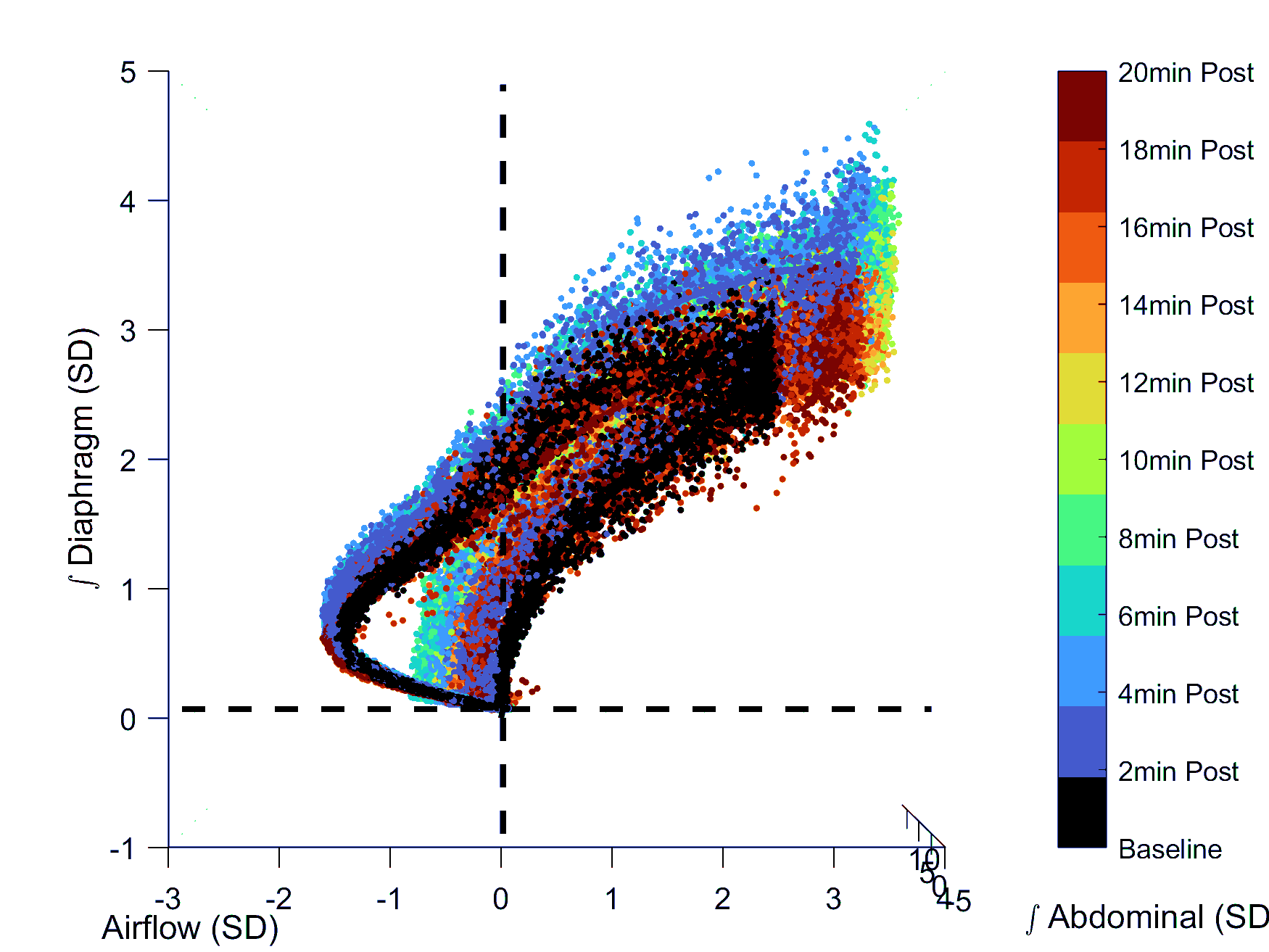
